# Uniform distribution of *35S* promoter-driven mDII auxin control sensor in leaf primordia

**DOI:** 10.1101/293753

**Authors:** Chunmei Guan, Fei Du, Yuling Jiao

## Abstract

The DII sensor has been an invaluable tool to map spatiotemporal auxin response and distribution in the model plant *Arabidopsis thaliana.* The DII sensor and mDII control sensor are driven by the widely used constitutive *35S* promoter. However, the reliability of DII sensor has recently been questioned (Bhatia and Heisler, 2018). Here we provide additional evidence to show that the mDII control sensor is indeed uniformly distributed in early leaf primordia, which echoes the original reports (Vernoux et al., 2011; Brunoud et al., 2012). We also use DII/mDII and the *PRS5A* promoter-driven R2D2 sensors to confirm asymmetric auxin signaling in early leaf primordia. On the other hand, we provide evidence that light penetration may lead to artifacts during whole-mount imaging.

## Main text

Auxin is a classical phytohormone that serves as a plant morphogenetic signal throughout the life cycle. Spatiotemporal distribution of auxin response is pivotal to understanding auxin’s role in pattern formation. DII-Venus is a recently developed auxin sensor, in which the fast maturing Venus fluorescent protein is fused in-frame to the auxin dependent degradation domain II of an Aux/IAA protein (Vernoux et al., 2011; Brunoud et al., 2012). It has been shown that the abundance of DII-Venus is dependent on auxin, the TIR1/AFBs co-receptors, and proteasome activities. In the original design, DII-Venus is driven by the widely used constitutive *35S* promoter. The control sensor, mDII-Venus, has a mutation in the domain II sequence of DII-Venus, thus lacking auxin-dependent degradation. The mDII-Venus control sensor is also driven by the same constitutive *35S* promoter. Comparing with mDII-Venus, reduced DII-Venus signal would indicate higher auxin. The DII-Venus sensor allows sensitive detection of auxin signaling at cellular resolution in different tissues, and has been widely used, including in the field of plant development. An alternative design, named R2D2, combines DII-Venus and mDII-tdTomato into a single transgene (Liao et al., 2015). In addition, the R2D2 uses the *RPS5A* promoter instead of the *35S* promoter.

The *35S*-driven mDII-Venus has uniform expression in the vegetative shoot apex, including the shoot apical meristem (SAM) and young leaf primordia (Fig. 2h in Brunoud et al., 2012; Supplementary Fig. 7b in Vernoux et al., 2011), making it possible to use the *35S*-driven DII-Venus to map auxin signaling. Compared with mDII-Venus, DII-Venus shows clearly stronger signal in the adaxial domain facing the SAM in the first two leaves five days after stratification (DAS) (Fig. 2f,h in Brunoud et al., 2012; Supplementary Fig. 7a in Vernoux et al., 2011). We observed similar patterns in early leaf primordia of older plants from 1-4 weeks after stratification (Qi et al., 2014; Guan et al., 2017), suggesting a transient abaxial-enrichment of auxin. These observations and the middle domain-enriched DR5 expression, together with genetics and molecular analysis, lead to a model, in which abaxial-enriched auxin, the adaxial-expressed auxin activator MONOPTEROS/AUXIN RESPONSE FACTOR5 (MP/ARF5), and abaxial-expressed ARF repressors together position auxin signaling in the middle domain (Guan et al., 2017). Auxin signaling activates *WUSCHEL-RELATED HOMEOBOX* (*WOX*) genes in the middle domain to enable leaf blade expansion.

In contrast to all above reports, Bhatia and Heisler very recently claimed that the *35S*-driven mDII-Venus control sensor also exhibits an adaxial-enrichment, which is similar to the *35S*-driven DII-Venus sensor, in the first two leaves of 3-4 DAS seedlings (Bhatia and Heisler, 2018). They further claimed that the *35S* promoter activity is in general strongly enriched in the adaxial domain than the abaxial domain, regardless of the reporter gene used.

In addition to previous mDII-Venus imaging results, this claim goes clearly against other imaging and genetic analysis data. For example, detailed imaging analysis clearly showed comparable activity of the *35S* promoter activity in adaxial and abaxial domains (Skopelitis et al., 2017). In addition, *35S*-driven expression of adaxial genes, such as *AS2* (Lin et al., 2003), results in adaxialized leaves, suggesting the *35S* promoter is active in the abaxial domain.

To understand if *35S*::*mDII-Venus* may have indeed a differential expression between the ad- and abaxial sides in the first pair of true leaves, we followed the sample stage and imaging setups of Bhatia and Heisler. Consistent with all previous reports in shoot apices of other stages (Vernoux et al., 2011; Brunoud et al., 2012; Wang et al., 2014), but different from Bhatia and Heisler, mDII-Venus showed clear uniform distribution throughout leaf primordia qualitatively and quantitatively (Figure 1 and S1). In very young leaf primordia, variability among plants and leaves is observed. In particular, the epidermis tends to have strong signal variability for mDII-Venus in 2 DAS seedlings (Figure 1A-F, S1, and also see Figure 5A and S3A). Nevertheless, there is no consistent enrichment for mDII-Venus in adaxial or abaxial domains. Albeit some signal variability, DII-Venus is adaxial enriched at the same stage (Figure 2A-F). At 3 DAS, the mDII-Venus signal is less variable (Figure 1G-L, S1G-R, see also Figure 5A, S3B and C), and the DII-Venus sensor was reproducibly adaxially enriched (Figure 2G-L), as shown before (Vernoux et al., 2011; Brunoud et al., 2012; Qi et al., 2014; Bhatia and Heisler, 2018). Bhatia and Heisler also used the R2D2 sensor to assess auxin signaling distribution in leaf primordia. We actually found slightly adaxial-enriched signals of the *pRPS5A*::*mDII-tdTomato* of R2D2 (Figure 3 and S2). However, the signal of *pRPS5A*::*DII-Venus* of R2D2 was more strongly adaxial-enriched. Ratiometric analysis of the two channels in R2D2 shows indeed adaxial-enriched of the DII-VENUS to mDII-VENUS ratio quantitatively (Figure 4). Taken together, both *35S*- and *RPS5A*-driven DII/mDII sensors indicate adaxial-enriched DII signals in 2-3 DAS seedlings, which is consistent with results obtained from leaf primordia of older plants.

**Figure 1.**
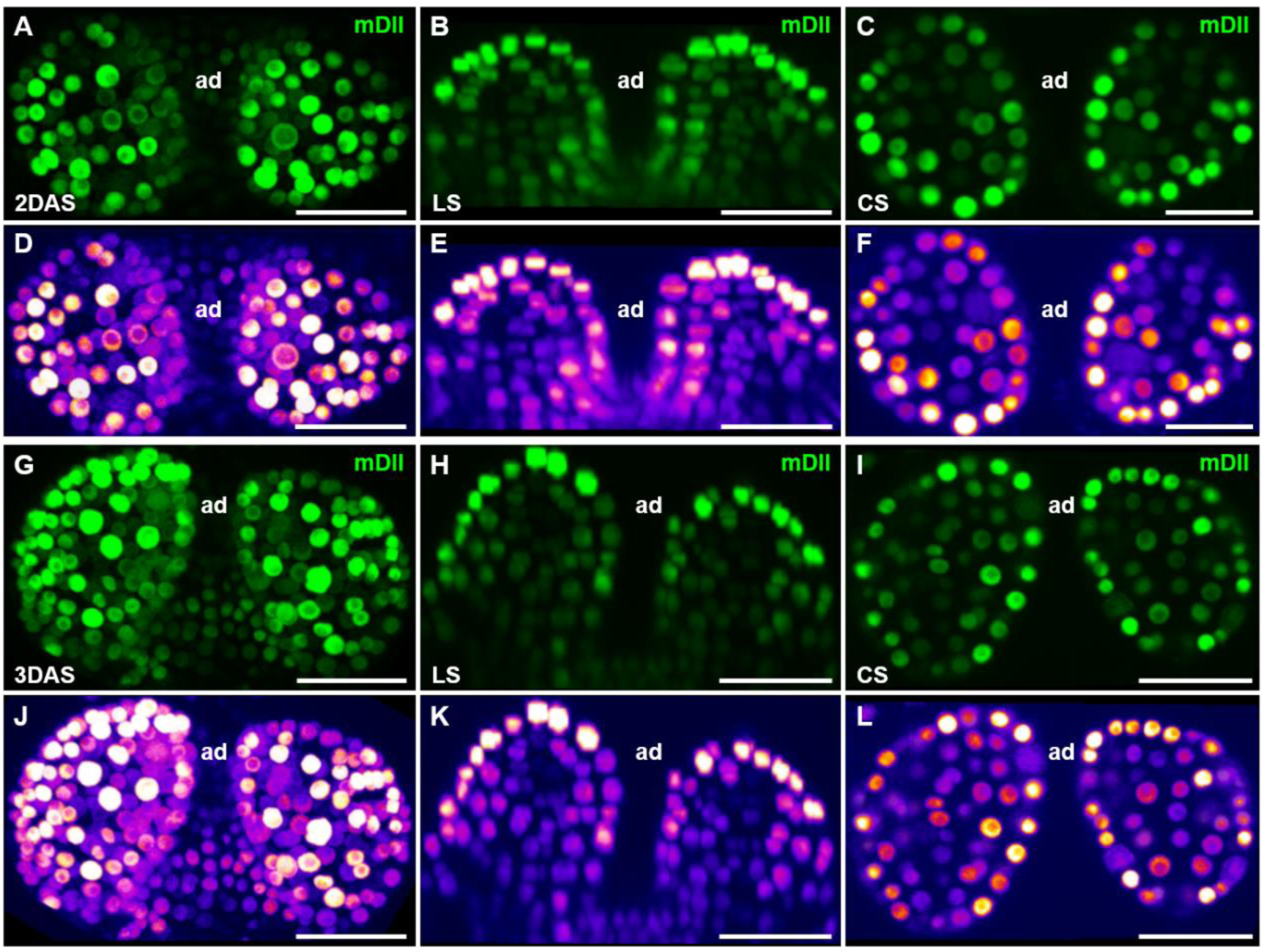
mDII signals in young Arabidopsis leaves. Optical sections through 2 DAS (**A** to **F**) and 3 DAS (**G** to **L**) shoot apices showing expression of mDII-Venus. For each panel, a confocal projection (**A** and **G**), a longitudinal section (LS, **B** and **H**), and a cross section (CS, **C** and **I**) are shown from left to right. (**D** to **H**, and **J** to **L**) Heatmaps of **A** to **C** and **G** to **I** showing the fluorescence intensity of mDII-Venus, respectively. ad, adaxial. Scale bars, 20 μm.

**Figure 2.**
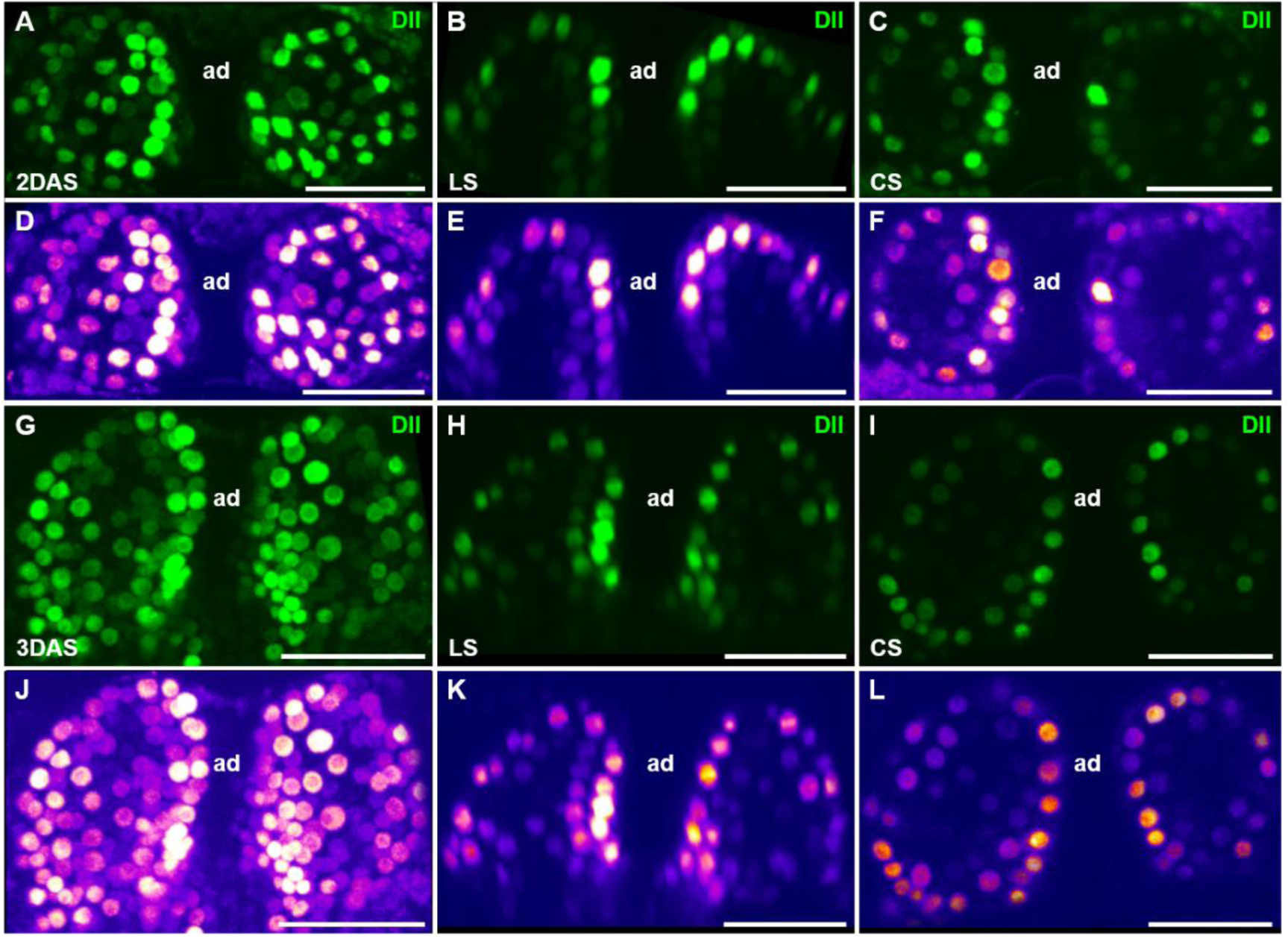
DII signals in young Arabidopsis leaves. Optical sections through 2 DAS (**A** to **F**) and 3 DAS (**G** to **L**) shoot apices showing expression of DII-Venus. For each panel, a confocal projection (**A** and **G**), a longitudinal section (LS, **B** and **H**), and a cross section (CS, **C** and **I**) are shown from left to right. (**D** to **H**, and **J** to **L**) Heatmaps of **A** to **C** and **G** to **I** showing the fluorescence intensity of DII-Venus, respectively. ad, adaxial. Scale bars, 20 μm.

**Figure 3.**
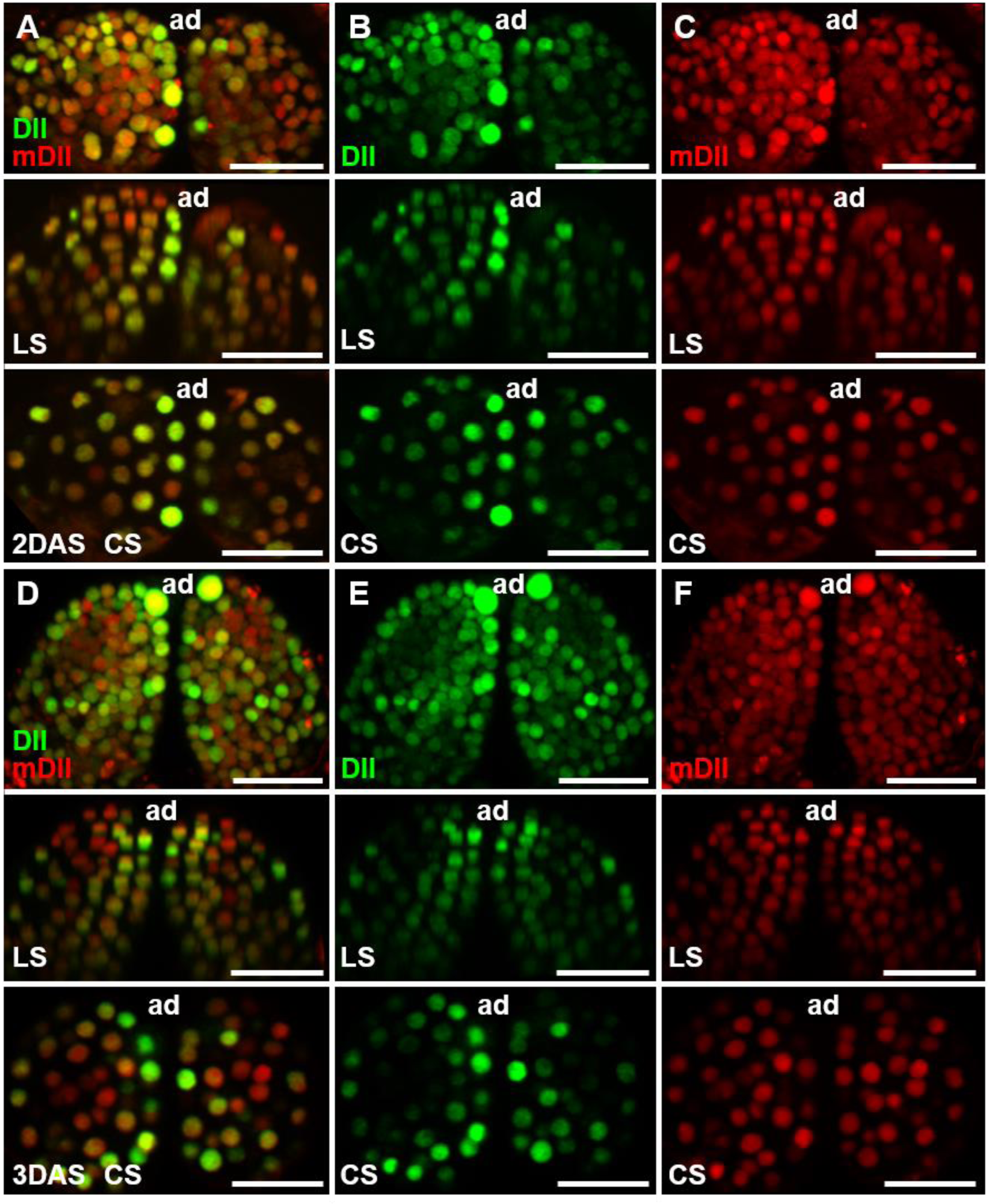
R2D2 signals in young leaves. Optical sections through 2 DAS (**A** to **C**) and 3 DAS (**D** to **F**) shoot apices showing expression of DII-Venus (green) and mDII-tdTomato (red). For each panel, a confocal projection, a longitudinal section (LS), and a cross section (CS) are shown from top to bottom. A top view is shown for the projection in (**A** to **C**), and a side view is shown for the projection in (**D** to **F**). Scale bars, 20 μm.

**Figure 4.**
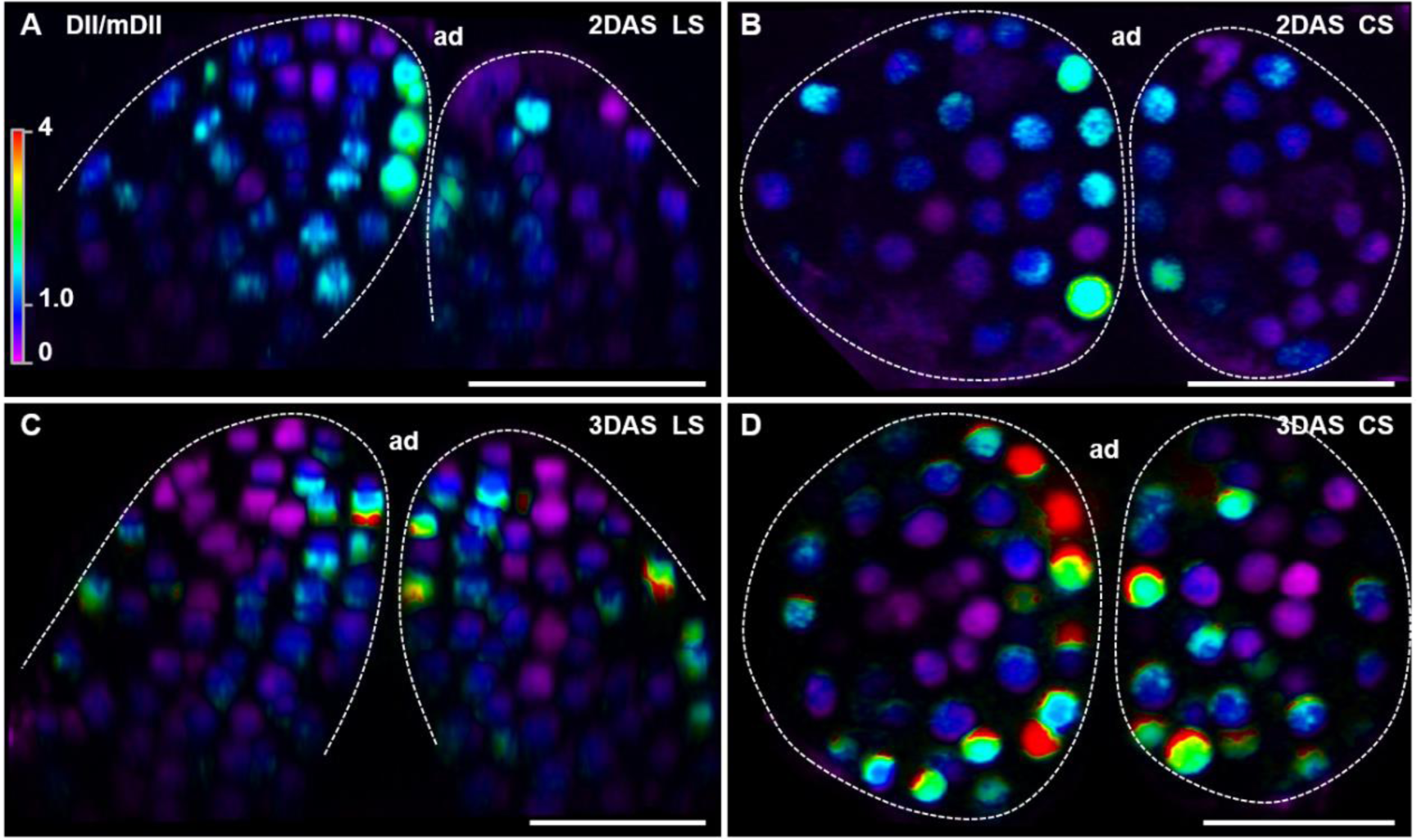
DII/mDII ratios of R2D2 in young leaves. Optical sections through 2 DAS (**A** and **B**) and 3 DAS (**C** and **D**) shoot apices showing DII-Venus/mDII-tdTomato signal ratios. Ratio calculations for DII/mDII were performed using Nikon instruments software NIS-Elements AR. Longitudinal sections (LS, **A** and **C**), and cross sections (CS, **B** and **D**) are shown. Scale bars, 20 μm.

**Figure 5.**
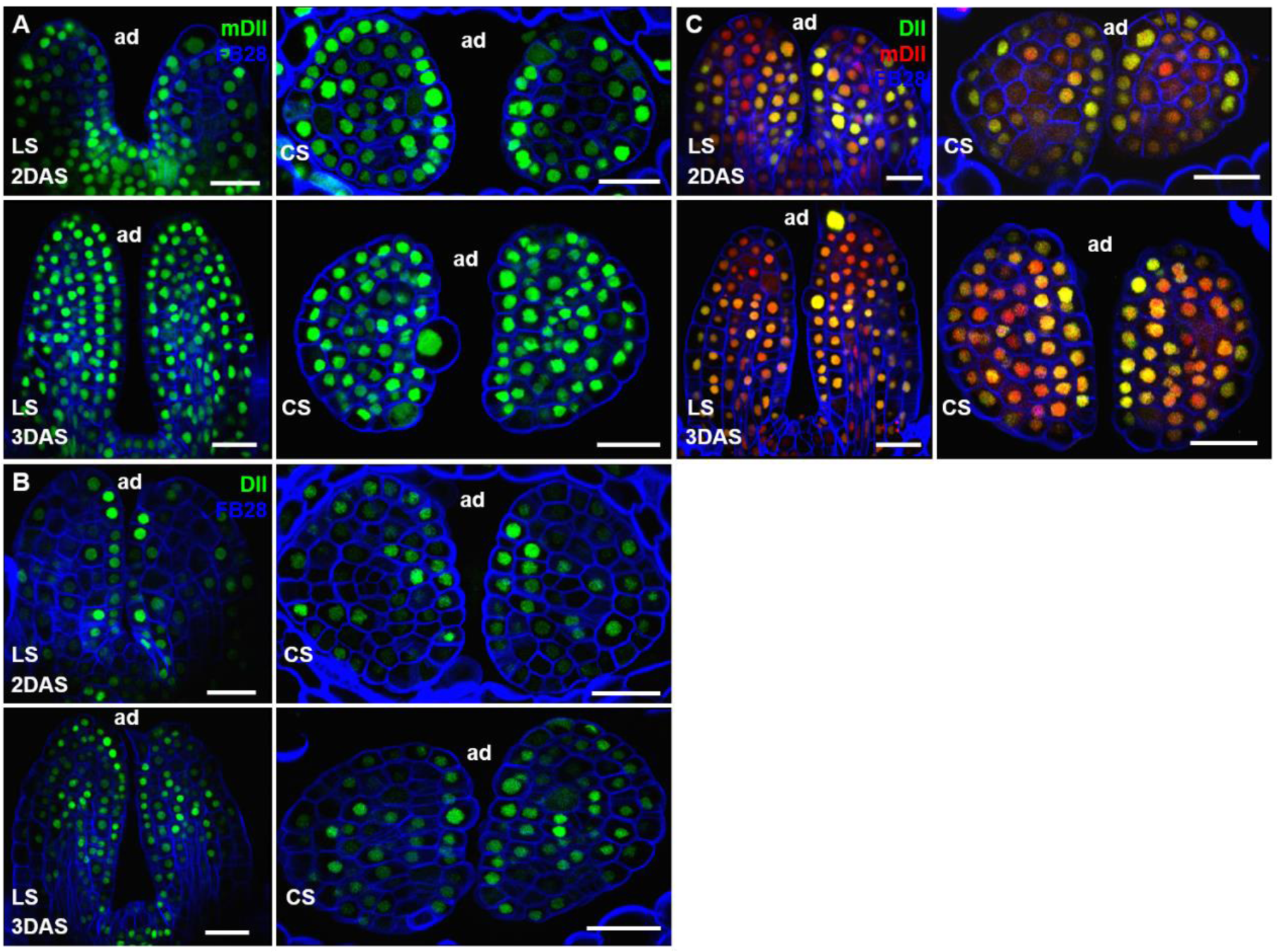
mDII and DII signals in microtome-based sections of young leaves. Thin sections through 2 DAS and 3 DAS shoot apices showing expression of mDII-Venus (**A**), DII-Venus (**B**), and R2D2 (**C**, red for mDII-tdTomato, and green for DII-Venus). Cell walls were stained with FB28 (blue). For each stage, a longitudinal section (LS), and a cross section (CS) are shown. Scale bars, 20 μm.

We reasoned that tissue shape may lead to biased signals when a whole-mount approach is used as Bhatia and Heisler did. It is widely known that fluorescence microscopy can only penetrate limited tissue depths. Several factors, including absorption by pigments and cytoplasmic components, and light scattering, may influence the tissue depth that can be penetrated. To avoid potential bias caused by signal penetration, we used mechanical sectioning to obtain high resolution images (Figure 5, S3 and S4). In vibrating microtome sections, it is evident that mDII-Venus and mDII-tdTomato have similar levels of signals in the adaxial and abaxial domains. In contrast, DII-Venus has adaxial-enriched signals. Thus, mechanical sectioning convincingly supports uniform activities of *35S* promoter in leaf adaxial and abaxial domains. Notably, the signal variability of mDII-Venus in 3 DAS seedlings is clearly lower in mechanical sections than in optical sections (compare Figures 5A and S3B-C with Figures 1G-L and S1G-R), indicating that whole-mount imaging may artificially alter signals.

Taken together, we used light sectioning in whole-mount samples, and mechanical sectioning to reanalyze *35S*::*mDII-Venus* distribution in young leaves of early seedlings. Our new analysis is consistent with previous imaging results (Vernoux et al., 2011; Brunoud et al., 2012; Wang et al., 2014). A uniform activity of the *35S* promoter is also supported by imaging other *35S*-driven fluorescent proteins, and by phenotypic analysis of *35S*-driven transgenic plants (Lin et al., 2003; Skopelitis et al., 2017). Thus, the asymmetric *35S*::*mDII-Venus* distribution reported by Bhatia and Heisler is in contrast to the results of our analysis. Adaxial-enriched DII-Venus is also consistent with additional genetic and imaging data. For example, leaves of *pMP*::*MPΔ*, which is active independent of auxin, have ectopic auxin signaling, *WOX* expression, and growth in the adaxial domain (Krogan and Berleth, 2012; Qi et al., 2014; Guan et al., 2017). Because *MP* expression encompass the adaxial domain (Rademacher et al., 2011; Qi et al., 2014; Guan et al., 2017), a low level of auxin is essential to avoid ectopic activation of *MP* downstream genes.

## Methods

### Plant material and growth conditions

The transgenic lines *p35S*::*DII*-*Venus* and *p35S*::*mDII*-*Venus* were obtained from Dr. Teva Vernoux, and the R2D2 reporter line was obtained from Dr. Dolf Weijers. Seeds were kept for 2 days on MS medium containing 1% sucrose in the dark at 4°C and then in the greenhouse under short-day-conditions for 2 or 3 days before imaging.

### Agarose gel sectioning

Agarose gel sectioning procedure was performed as previously described (Skopelitis et al., 2017) with minor modifications. Seedlings grown under short-day-conditions for 2 or 3 days were collected into freshly prepared fixative solution containing 4% paraformaldehyde and 0.015% Tween-20 in 1×PBS (pH = 7.0). Vacuum infiltration at -0.075 MPa was performed twice for 10 min each time. Seedlings were washed three times (10 min per time) in 1 × PBS and embedded into 6% low melting agarose (Promega). We obtained 40-50 μm transverse or longitudinal sections using a VT1000S vibratome (Leica). Agarose sections were stained with 0.01% Fluorescent Brightener 28 (FB28) (Sigma-Aldrich) dissolved in 1×PBS for 20 min in darkness, followed by three times wash with 1×PBS (5 min per time). Sections were mounted in 90% glycerol in 1×PBS for confocal microscopy.

### Confocal imaging and data analysis

For live imaging, seedlings grown under short-day-conditions for 2 or 3 days were transferred to the dissecting medium (3% agarose). Cotyledons were carefully removed under stereomicroscope (Nikon, SMZ18). Live imaging was performed using 60× water dipping lens. Imaging of vibratome sections was performed using 60× oil immersion lens. Confocal images were taken with a Nikon A1 confocal microscope. Excitation and detection windows setups for Venus and tdTomato were as described (Qi et al., 2014). To image Venus, a 514 nm laser excitation and a 524-550 nm band-pass filter emission was used. To image tdTomato, a 561 nm laser was used for excitation and a 570-620 nm band-pass filter was used for detection. To image FB28, a 405 nm laser excitation and a 425-475 nm band-pass emission filter were used. DII/mDII ratio calculation was performed using Nikon NIS-Elements AR.

## Acknowledgements

We thank Dr. Teva Vernoux for the DII and mDII lines, and Dr. Dolf Weijers for the R2D2 line. We also thank Drs. Teva Vernoux and Ying Wang for critical comments on the manuscript.

---

**Figure S1.**
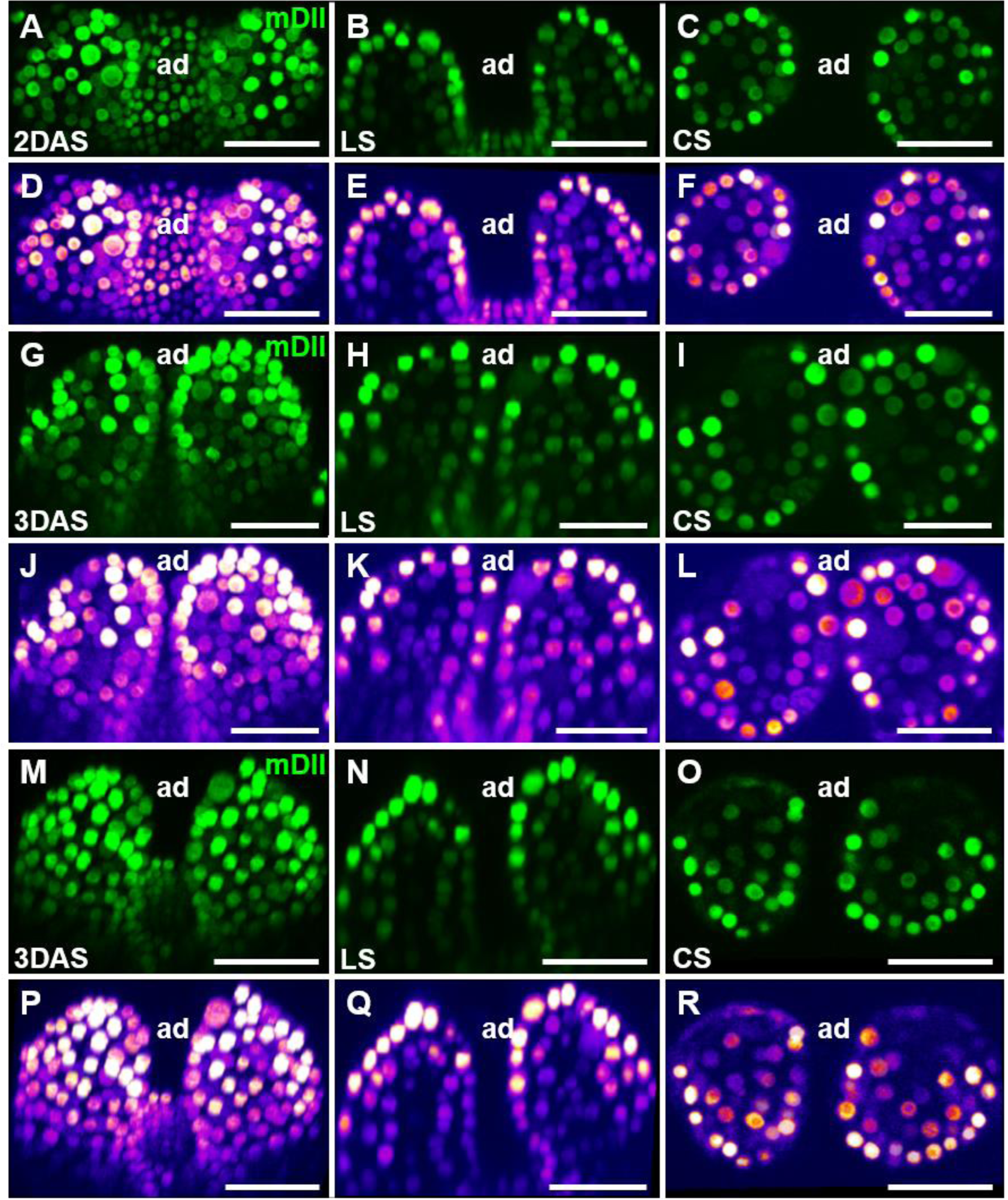
Additional examples of mDII signals in young Arabidopsis leaves. Optical sections through 2 DAS (**A** to **F**) and 3 DAS (**G** to **R**) shoot apices showing expression of mDII-Venus. For each panel, a confocal projection, a longitudinal section (LS), and a cross section (CS) are shown from left to right. Heatmaps showing the fluorescence intensity of mDII-Venus for each panel are shown below. ad, adaxial. Scale bars, 20 μm.

**Figure S2.**
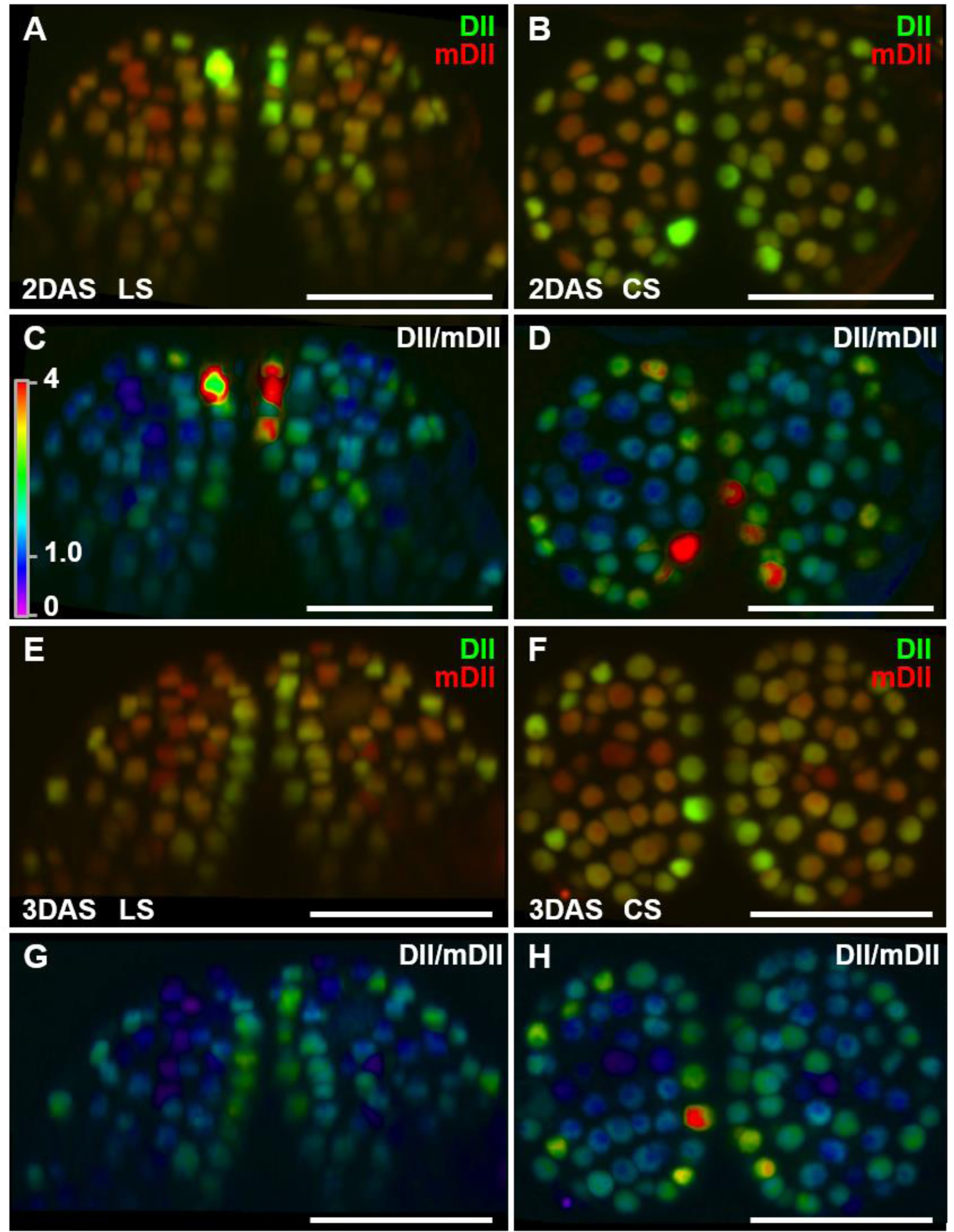
Additional examples of R2D2 signals in young Arabidopsis leaves. Optical sections through 2 DAS (**A** to **D**) and 3 DAS (**E** to **H**) shoot apices showing expression of DII-Venus (green) and mDII-tdTomato (red). For each panel, a longitudinal section (LS) and a cross section (CS) are shown from left to right. DII-Venus/mDII-tdTomato signal ratios for each panel are shown below. Scale bars, 20 μm.

**Figure S3.**
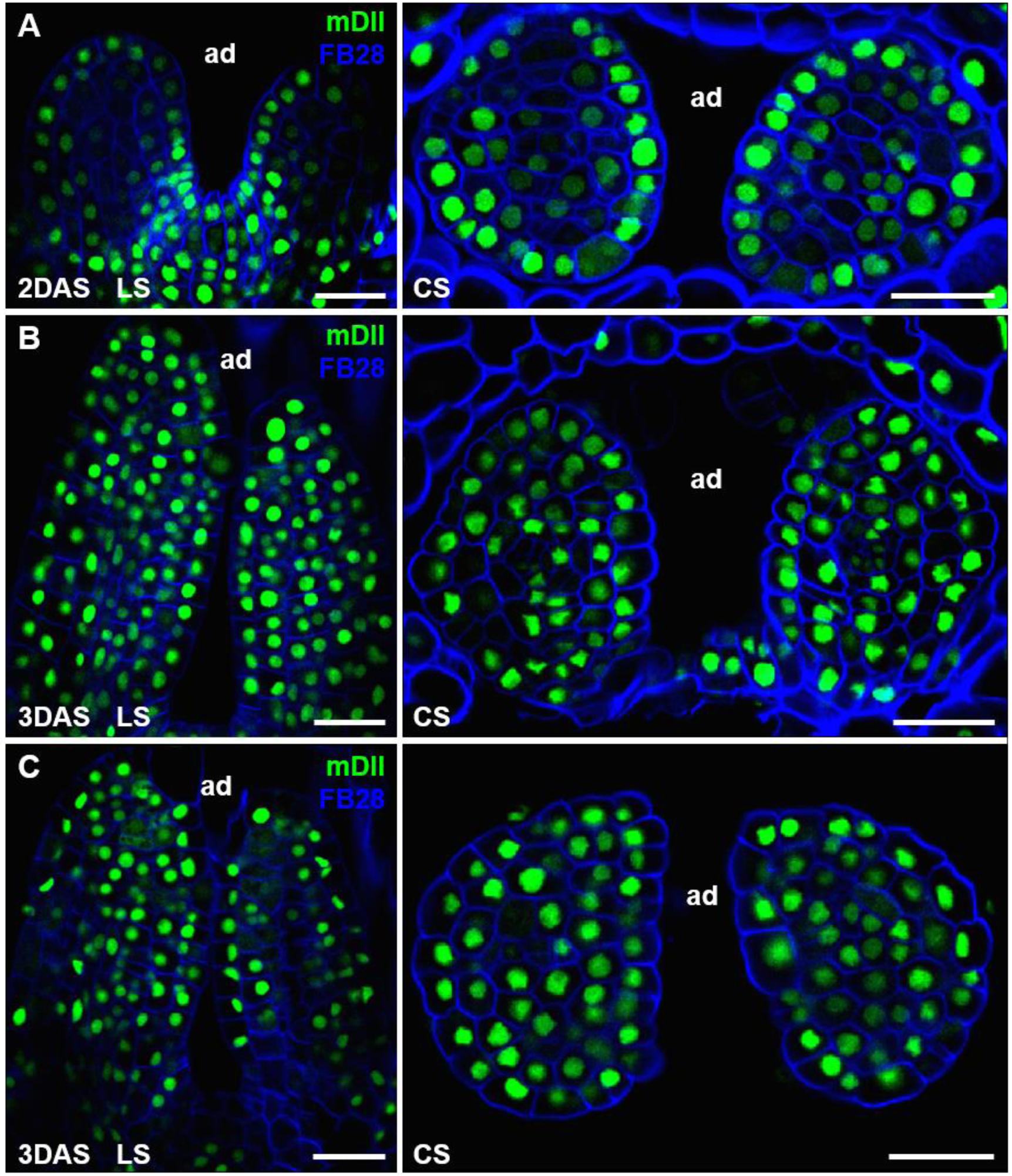
Additional examples of mDII signals in microtome-based sections of young leaves. Thin sections through 2 DAS (**A**) and 3 DAS (**B** and **C**) shoot apices showing expression of mDII-Venus (green). Cell walls were stained with FB28 (blue). For each stage, a longitudinal section (LS), and a cross section (CS) are shown. Scale bars, 20 μm.

**Figure S4.**
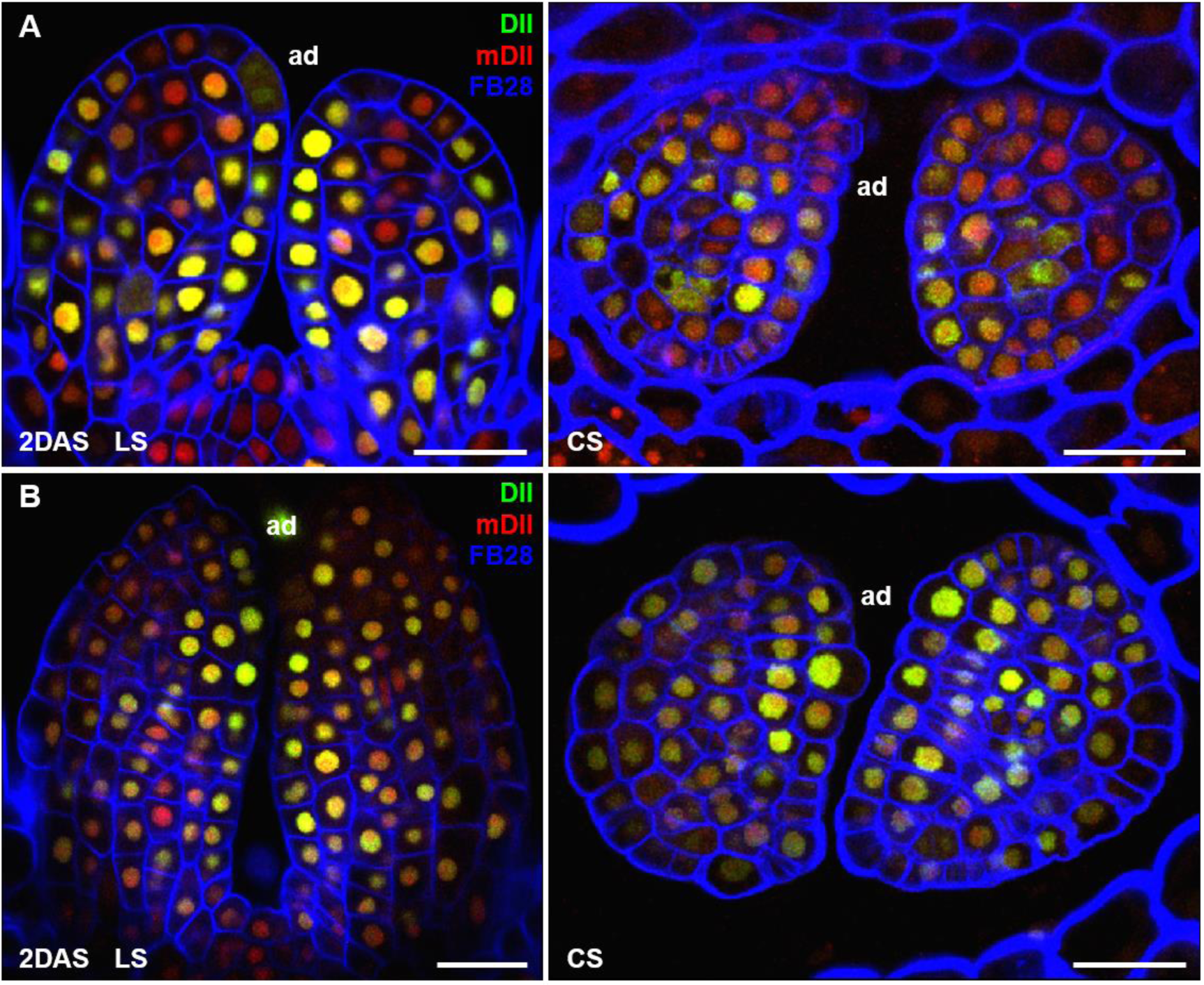
Additional examples of R2D2 signals in microtome-based sections of young leaves. Thin sections through 2 DAS (**A**) and 3 DAS (**B**) shoot apices showing expression of R2D2 (red for mDII-tdTomato, and green for DII-Venus). Cell walls were stained with FB28 (blue). For each stage, a longitudinal section (LS), and a cross section (CS) are shown. Scale bars, 20 μm.

## References

Bhatia, N., and Heisler, M.G. (2018). Auxin is not asymmetrically distributed in initiating Arabidopsis leaves bioRxiv, 10.1101/284554.

Brunoud, G., Wells, D.M., Oliva, M., Larrieu, A., Mirabet, V., Burrow, A.H., Beeckman, T., Kepinski, S., Traas, J., Bennett, M.J., and Vernoux, T. (2012). A novel sensor to map auxin response and distribution at high spatio-temporal resolution. Nature 482, 103–106.

Guan, C., Wu, B., Yu, T., Wang, Q., Krogan, N.T., Liu, X., and Jiao, Y. (2017). Spatial auxin signaling controls leaf flattening in *Arabidopsis*. Curr Biol 27, 2940–2950.

Krogan, N.T., and Berleth, T. (2012). A dominant mutation reveals asymmetry in MP/ARF5 function along the adaxial-abaxial axis of shoot lateral organs. Plant Signal Behav 7, 940–943.

Liao, C.Y., Smet, W., Brunoud, G., Yoshida, S., Vernoux, T., and Weijers, D. (2015). Reporters for sensitive and quantitative measurement of auxin response. Nat Methods 12, 207–210.

Lin, W.-C., Shuai, B., and Springer, P.S. (2003). The *Arabidopsis* LATERAL ORGAN BOUNDARIES-domain gene *ASYMMETRIC LEAVES2* functions in the repression of KNOX gene expression and in adaxial-abaxial patterning. Plant Cell 15, 2241–2252.

Qi, J., Wang, Y., Yu, T., Cunha, A., Wu, B., Vernoux, T., Meyerowitz, E., and Jiao, Y. (2014). Auxin depletion from leaf primordia contributes to organ patterning. Proc Natl Acad Sci U S A 111, 18769–18774.

Rademacher, E.H., Moller, B., Lokerse, A.S., Llavata-Peris, C.I., van den Berg, W., and Weijers, D. (2011). A cellular expression map of the *Arabidopsis AUXIN RESPONSE FACTOR* gene family. Plant J 68, 597–606.

Skopelitis, D.S., Benkovics, A.H., Husbands, A.Y., and Timmermans, M.C.P. (2017). Boundary formation through a direct threshold-based readout of mobile small RNA gradients. Dev Cell 43, 265–273.

Vernoux, T., Brunoud, G., Farcot, E., Morin, V., Van den Daele, H., Legrand, J., Oliva, M., Das, P., Larrieu, A., Wells, D., Guedon, Y., Armitage, L., Picard, F., Guyomarc’h, S., Cellier, C., Parry, G., Koumproglou, R., Doonan, J.H., Estelle, M., Godin, C., Kepinski, S., Bennett, M., De Veylder, L., and Traas, J. (2011). The auxin signalling network translates dynamic input into robust patterning at the shoot apex. Mol Syst Biol 7, 508.

Wang, Y., Wang, J., Shi, B., Yu, T., Qi, J., Meyerowitz, E.M., and Jiao, Y. (2014). The stem cell niche in leaf axils is established by auxin and cytokinin in *Arabidopsis*. Plant Cell 26, 2055–2067.

